# The relationship between confidence and gaze-at-nothing oculomotor dynamics during decision-making

**DOI:** 10.1101/2024.08.29.610272

**Authors:** Ignasi Cos, Gizem Senel, Pedro E. Maldonado, Rubén Moreno-Bote

## Abstract

How does confidence relate to oculomotor dynamics during decision-making? Do oculomotor dynamics reflect deliberation and the buildup of confidence in the absence of visual stimuli? Here we examine the hypothesis that working memory, deliberation, and confidence warp oculomotor dynamics, both in the presence and absence of visual stimuli. We analyzed oculomotor dynamics in a decision-making task in which participants were provided with an abstract context in which to make the decision, and two similar option images which became eventually invisible. We show that fixations between the empty locations in which the images were formerly shown continued after the options disappeared, consistently with a sustained deliberative process facilitated by oculomotor dynamics. Both, oculomotor dynamics and decision patterns remained unchanged regardless of whether the stimuli were visible. Furthermore, our analyses show that the number of alternative fixations between stimuli correlated negatively with the confidence reported after each decision, while the observation time of the selected target correlated positively. Given that decisions in our experimental paradigm are reported in the absence of the stimuli, this suggests a relationship between evidence retrieval from working memory, confidence gathering and oculomotor dynamics. Finally, we performed a model comparison based on predictions from drift-diffusion models to assess the relationship between sequential fixations between images, deliberation and confidence gathering, and the ensuing choice. These analyses supported confidence as a contributing cognitive process encompassing serial evidence-gathering and parallel option consideration during decision-making.

**One-Sentence Summary:** The dynamics of oculomotor dynamics between absent stimuli are related with the participant’s confidence during value-based decision-making.

## INTRODUCTION

Decision-making entails evidence gathering and commitment to a particular option when the evidence is deemed sufficient^1–3^. As evidence is collected over time, individuals develop a subjective sense of confidence, which they may use to determine when to terminate the gathering of evidence and commit to an option^4,5^. Confidence estimation has been described as a cognitive process monitoring decision-making^5–9^, providing an internal metric of the reliability of the evidence and the trustworthiness of the choices^4,8,10^. It participates in all aspects of decision-making, from perception to the execution of motor responses^11^, and has been used as a proximal measure for cognitive performance^12^ and choice accuracy^5,13,14^. While much research has focused exclusively on the perceptual aspect of decision making, it have been long acknowledged that motor aspects of the choices also play a significant role in option selection^15–18^. Nonetheless, the specifics of how sensorimotor aspects of decision-making contribute to the confidence estimation remains unsettled^19^.

In visually-guided decision-making, evidence is gathered via sequential sampling/oculomotor fixations over the details of each image stimulus^20^, to be transferred from sensory to working memory representations in a process modulated by evidence coherence/complexity^21^. Consistent with this, observation times and the number of saccades typically increase with more difficult perceptual discriminations. This strongly suggests that option analysis and deliberation^5^, along with its underlying subjective sense of confidence, participate of this process. In line with this, oculomotor processes have been shown to be predictive of the likely choice to follow, typically the stimuli fixated longer and/or more frequently^22^. However, the question remains whether deliberation and oculomotor control are segregated processes – as suggested by sequential sampling models^1,3,12^ and previous studies in visual decision-making^23^ –, or whether they are intrinsically related during decision-making. If the latter is the case, we should expect that oculomotor dynamics contribute to option assessment and working memory deliberation even when there is no sensory evidence gathered.

To address these questions, and by contrast to prior experimental paradigms using perceptual decisions, we needed a task in which the participants had to make complex judgements in response to abstract goals, which did not directly depend on the mere sensory properties of the stimuli presented. Thus, we designed a context-dependent task in which image options offer similar utility, at least a priori, and eventually disappear during the course of the trial, by extending from the *look-at-nothing* effect paradigm^24^. This is a visuo-motor effect consisting of looking at empty locations where images were formerly presented to aid with memory retrieval. This design is appropriate for three reasons: first, eye movements were *not* constrained to fixation points; second, the absence of stimuli during specific phases of the trial ensured that eye movements could be decoupled from the gathering of visual evidence from external stimuli and related with internal deliberative processes in working memory; third, to promote deliberation, the contexts in which the decisions were made were unusual and the images presented offered options to have the similar value in those contexts (see list of items and contexts in Supplemental).

In brief, in our novel task participants were presented with a text describing the context in which the decision of that trial would unfold (Figure 1; e.g., What would you rather have a nightmare with?). This was followed by the presentation of two images (an evil clown vs. a witch) representing the two options within the referred context. Participants were instructed to choose one of both options based on the option’s subjective appropriateness for the given context. Importantly, both images disappeared after a Stimulus-On presentation period --- indicating the GO signal, leaving behind two frames to indicate their former locations, and starting the Stimulus-Off interval during which the choice could be reported by pressing the right/left mouse button. Oculomotor movements were recorded both during stimuli presentation (look-at-something) and when stimuli were absent (looking-at-nothing). We show that oculomotor movements between empty locations continued after the stimuli disappeared, consistent with a sustained deliberative process facilitated by visuo-motor dynamics, which alternate focus between working memory representations. We found that the longer the observation of a specific empty location, the higher the probability of choosing the image formerly shown in that area. Furthermore, we also found that the higher the frequency of visual alternations between stimuli or their empty locations, the lower the confidence reported after the choice. Along with a model comparison based on predictions from parallel or sequential drift-diffusion models, our results are strongly consistent with working memory, deliberation and confidence to be processes intrinsically related to oculomotor dynamics, regardless of the presence/absence of visual stimuli.

**Figure 1.**
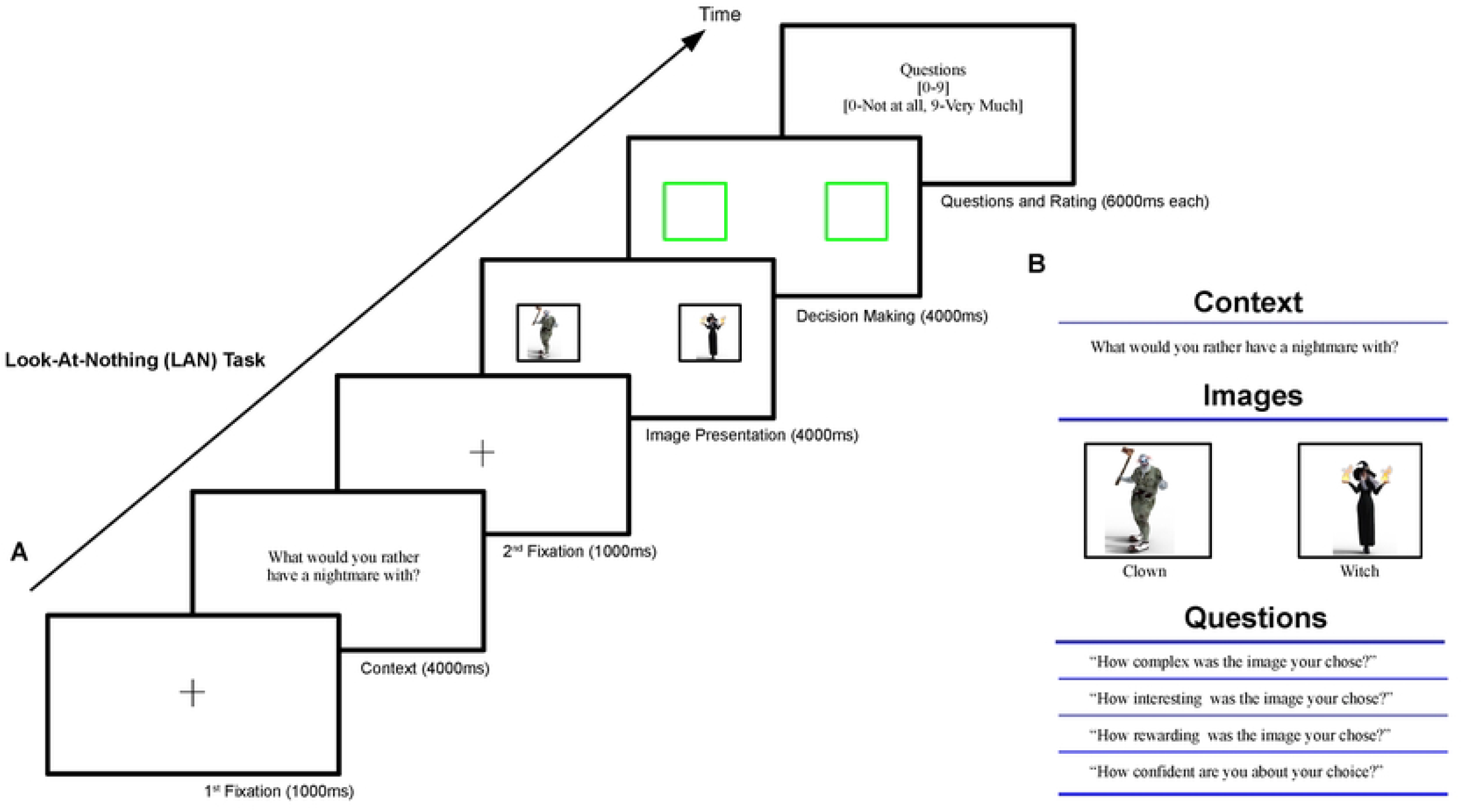
A context-dependent binary decision-making paradigm to assess visual fixation patterns during look-at-something and look-at-nothing in a deliberated decision. **A**. Time-course of a typical trial of the LAN experiment. Each trial starts with a white screen with a fixation crosswire in its center. After 1s the crosswire disappears, and a brief text describing of the context of the decision is shown. 4s later the context disappears, and a second fixation crosswire is shown in the center of the screen. 1s after that, the fixation crosswire disappears, and both images representing the options to select upon are shown on the right and left of the screen, initiating the Stimulus-On interval. The GO signal for the participant to report his/her choice was given 4s later, when both images disappear while leaving their frames behind (in green). This starts the Stimulus-Off interval. These frames disappear 4s later, initiating a sequence of four questions about the choice, lasting 6s each: how complex was the decision, how interesting was the choice, how rewarding was the choice, and how confident were you about your decision. The trial ends with an empty white screen lasting 1s, acting as an inter-trial interval (ITI). **B**. Schematic of the sequence for a typical trial. We first provided the context in which both options were to be gauged. Both images represent the options to select upon. The four questions listed below characterize four aspects of each decision participants are asked about: complexity, interest, reward and confidence.

## RESULTS

In the main experimental task, which we called *Look-At-Nothing* (LAN), participants were presented with an abstract context in which to make a choice, e.g. what to take to a deserted island, what would you rather have a nightmare with (see Supplemental Material for the full list of contexts). The context was followed by two images (Stimulus-On interval), on the right and left sides of the screen (See Methods; Figure 1), showing two options to select upon. Options were chosen so at to represent, at least a priori, options with similar utility. In the main tasks, the images disappeared after 4s, indicating the GO signal and starting the Stimulus-Off interval, in which two green empty frames replaced the images at their former location. From that time onwards, the participants could report their choice of option/image they deemed most appropriate given the context by pressing one of two keyboard keys (see Methods). To purposely promote deliberation, we used unfamiliar contexts and options of comparable utility (see Suppl. Material and repository: https://www.kaggle.com/datasets/novecentous/lanimages/data). After reporting their choice, the participants were required to quantify cognitive factors related to their decision with a number ranging from a minimum of zero to a maximum of ten: the confidence in their choice, the complexity, interest, and reward provided by the options presented (Figure 1B). The participant’s gazing trajectories were captured throughout each trial by means of an EyeTribe (EyeTribe Inc, DK) oculometer (see Methods). To complement the primary LAN experiment, we performed three additional controls: a baseline Look-At-Something (LAS) experiment, in which both options were always visible to control for the influence of absent stimuli; a Motor Reversal (rLAN) experiment, in which the reporting action (Right|Left) and the side of the option presented were reversed, to control for the potential influence of the motor response on the choice; and a Short Time Observation (sLAN) experiment, with a reduced Images-On interval (from 4 to 2s), to control for the possibility that a commitment could have been reached before the GO signal (despite task difficulty and the fact that options exhibit a similar utility therein).

As metrics to characterize the unfolding of the decision-making process at each trial, we computed the time spent looking at each image during the Images-On interval or at the empty frames during the Images-Off interval. We also counted the number of back-and-forth saccades between images and/or their empty locations. We first anticipated that having complex, unfamiliar contexts and options with a-priori similar value would induce significant deliberation and, consequently, prolonged decisions. This was confirmed by the long decision times (DT; from the GO signal and after a 4s Stimulus-On interval for the LAN, LAS and rLAN experiments, and after a 2s Stimulus-On interval in the sLAN experiment). Figure 2A shows the distributions of DTs for the LAN: Avg. 1404.94ms, SE 271.46ms; LAS: Avg. 1372.60ms, SE 216.45ms; rLAN: Avg. 1637.84ms, SE 477.95m; sLAN: Avg. 1395.40ms, SE 617.67ms. We found no statistical differences between the distributions of decision times of the LAN vs. LAS experiments (Figure 2B; t-test, t=0.70, CI=[-0.10, 0.22], df=29, P=0.48; Bayes Factor (null hypothesis)=5.55). Overall, this suggests that, all other factors being equal, deliberation in the LAN and LAS conditions extends over a similar time duration, thus suggesting a negligible contribution of the stimuli persistence to the decision during the decision phase. Likewise, we found no statistical differences between the LAN and sLAN conditions (t-test, t=1.533, CI=[-0.07 0.53], df=32, P=0.13; BF(null hypothesis)=2.46). By contrast, we found a marginally non-significant statistical difference when we reversed the laterality of motor responses (LAN vs. rLAN; t-test, t=-1.73, CI=[-0.36 0.029], df=32, P=0.09; BF(null hypothesis)=1.86), suggesting that reversing the laterality of the response required some inhibition.

**Figure 2.**
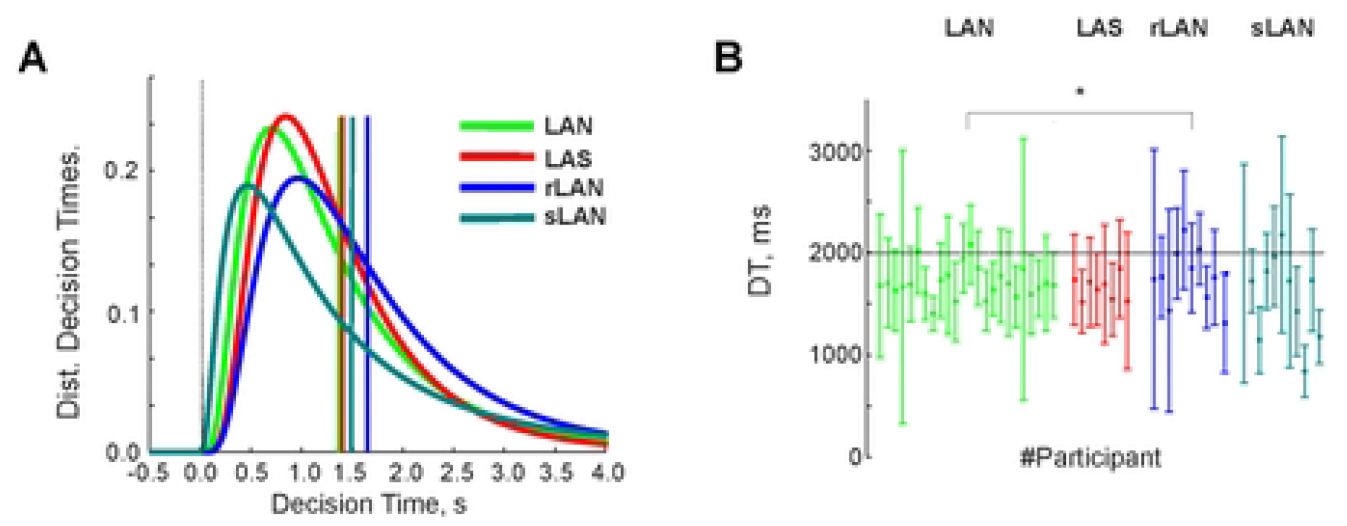
Decisions times between the look-at-something (LAS) and look-at-nothing (LAN) conditions do not differ. **A**. Histogram of Decision Times (DTs) for experiments LAN (red), LAS (green), rLAN (blue) and sLAN (turquoise). **B**. Mean and standard deviations of DTs for each participant and experiment LAN (green), LAS (red), rLAN (blue), sLAN (turquoise). All distributions were statistically undistinguishable except for the rLAN, which exhibited a further delay than the others (KS-test, LAN vs. rLAN, P=0.024).

In accord with previous literature^22^, we next hypothesized that the longer an option is observed, the more likely becomes its selection, with the novelty that here the image-stimuli were absent from the screen during the Stimulus-Off interval. We first tested during the Stimulus-On by calculating the probability of choosing the image on the right side of the screen (P_R_) as a function of the logarithmic ratio of times spent looking at the right over the left option for the LAN, LAS, rLAN, and sLAN experiments. In the LAN, LAS and sLAN, the P_R_ exhibits a steep increase as the proportion of observation time tilts in favor of the image on the right during *Stimulus-On* (LAN; Figure 3A & 3C; t-test across the participants’ slope β_1_ parameters, t=5.13, CI=[0.51, 1.22], df=21, P=4.41E-5; LAS; Figure 3F & 3H; t=4.69, CI=[0.54, 1.59], df=8, P=0.0016; sLAN; Suppl. Figure 1F & 1H, t=4.41, CI=[0.73 2.24], df=10, P=0.0013). The steep increase of P_R_ during the rLAN experiment is marginally non-significant, although consistent with the motor reversal between observation and reporting sides, in favor of the image on the left (Suppl. Figure 1A & 1C, t=-2.30, CI=[-1.68, 0.0017], df=8, P=0.050). Although this was expected while the option images were visible (Stimulus-On)^25,26^, we extended our analysis to the Stimulus-Off interval, during which the images were absent. Our analyses show that the effect persisted as well in the LAN experiment (Figure 3B & 3D; t-test across slope β_1_ parameters, t=3.60, CI=[2.20, 8.22], df=21, P=0.0017), in the sLAN experiment (Suppl. Figure 1G & 1I; t=5.03, CI=[0.59, 1.53], df=10, P=5.17E-4), in the rLAN experiment (Suppl. Figure 1B & 1D; t=-3.60, CI=[-1.29, -0.28], df=8, P=0.0069), and during the Stimulus-On interval after the GO signal for the LAS experiment (Figure 3G & 3I; t=2.79, CI=[0.81, 8.43], P=0.023). Most importantly, being all other conditions equal, we found no statistical difference between the LAN and LAS slope distributions neither during the Stimulus-On Observation Interval (t=0.64, CI=[-0.42,0.82], df=29, P=0.52; Bayes Factor(null hypothesis)=0.42), nor after the GO signal (Stimulus-Off LAN vs. LAS experiment (t=-0.0131, CI=[-0.25, 0.24], df=16, P=0.98; Bayes Factor (null hypothesis)=0.41). This is consistent with the contribution of sensory stimuli to deliberation during the decision phase is negligible.

**Figure 3.**
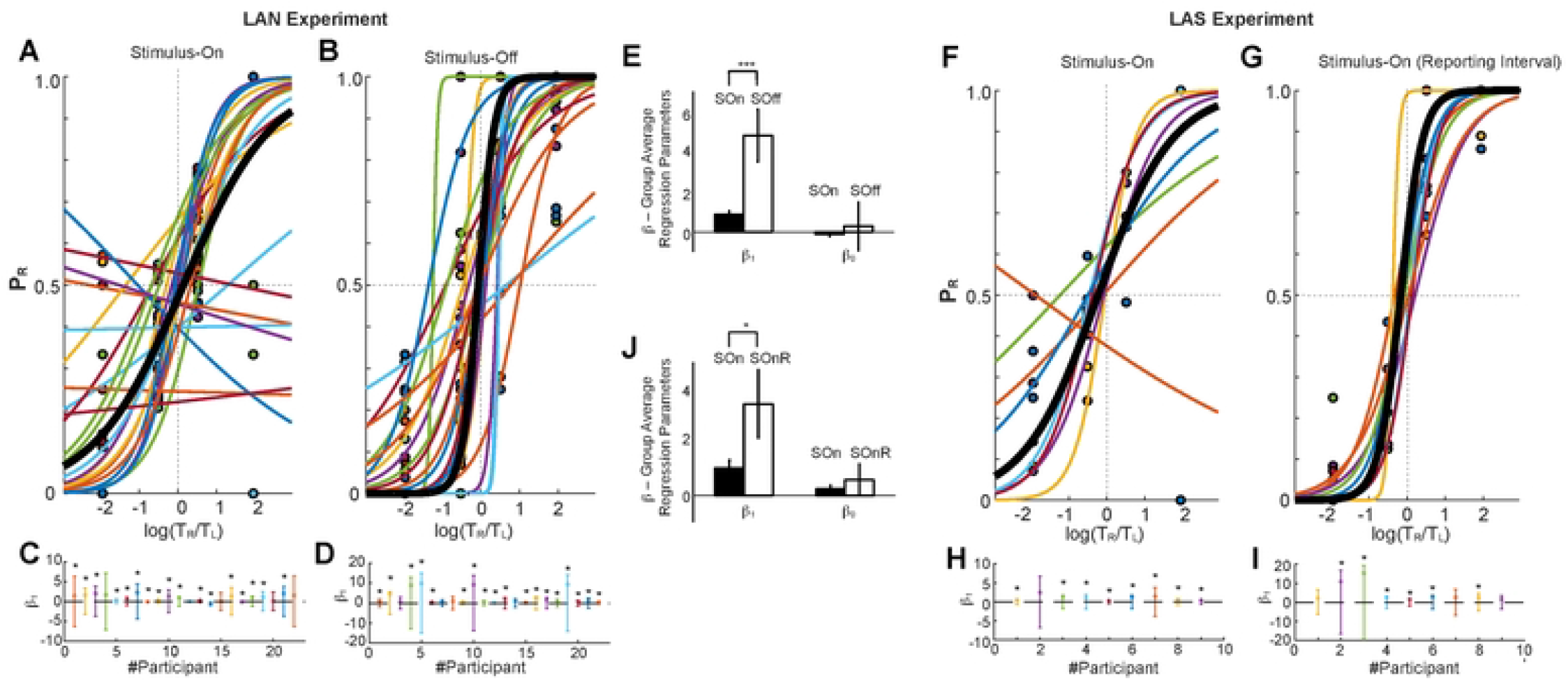
Looking times predict choices when visual options are visible and when they are invisible in a similar manner. **A-B**. LAN Experiment Analyses. Estimate of the probability of selecting the Right image as a function of the ratio of times spent looking Right over looking Left images during the Stimulus-On (A) and during Stimulus-Off (B) intervals. Sigmoidal regression fit for each participant and grand average (in black). **C-D**. Participants’ sigmoid slope parameter (β_1_) per participant during the Stimulus-On (C) and Stimulus-Off (D) intervals of the LAN experiment. Statistical significance was reported (* P<0.05). **E**. Comparison of the distribution of the participants’ sigmoid slope (β_1_) and intercept (β_0_) parameters between the Stimulus-On (SOn) and Stimlus-Off (SOff) intervals. **F-J**. Same as **A-E**, but for the LAS experiment. Note that in the LAN task there was no Stimulus-Off interval after the GO signal, but rather a second Stimulus-On interval, as both images were retained on the screen while the decision was reported.

We also confirmed that the β_1_ slope parameters obtained from the P_R_ fits during the *Stimulus-Off* interval were significantly steeper than those during the *Stimulus-On* interval in the LAN experiment (Figure 3E; t-test, t=-2.97, CI=[-7.27,-1.39], df=42, P=0.0048), and, marginally, in the LAS Experiment (Figure 3J, Stimulus-On vs. Stimulus-On after GO; t-test, t=-2.13, CI=[-7.08, -0.014], df=16, P=0.049; Figure 3J). This is consistent with a continuous decision-making process forming over time, starting during the Stimulus-On interval and extending onto the Stimulus-Off/Stimulus-On interval after GO. Importantly, this also consistent with choice commitment being formed after the GO signal. The increasingly steeper slopes with time are also visible in the case of the rLAN task (Suppl. Figure 1A-D; t-test, t=-0.12, CI=[-0.95, 0.85], df=16, P=0.91; Suppl. Figure 1E) and sLAN task (Suppl. Figure F-I; t-test, t=1.08, CI=[-0.39, 1.26], df=20, P=0.29; Suppl. Figure 1J), although these effects were not statistically significant.

In so far, we have shown that oculomotor dynamics are consistent with a long deliberation that extends into the decision-reporting phase, both in tasks in which the visual stimuli to decide upon are present (LAS) or absent (LAN). However, a relevant question is whether oculomotor dynamics are also related with confidence during deliberation. If that were the case, we first expected that longer decision times would accompany more difficult and thus less confident decisions. More importantly, we predicted that looking at an image (or its empty frame) relative to the other image for a long time, would relate to a higher confidence in the decision, while more balanced observation times or more frequent alternations between images would be indicative of a lower confidence (see Methods).

Consistent with the first prediction, we found a strong negative correlation between the reported confidence index and the decision time in three out of four experiments (LAN, Figure 4C, 4G, Avg. Correlation Index=-0.26 (±0.032 SE), t-test, t-stat=-8.58, CI = [-0.33, -0.20], df =23, P=1.26E-8; LAS, Figure 4F-G; Avg. Correlation Index=-0.10 (±0.062 SE), t-test, t-stat=-3.40, CI = [-0.35, 0.0090], df =7, P=0.0059; rLAN, Suppl. Figure 1K & Figure 4G, Avg. Correlation Index: -0.27 (±0.042 SE), t-test, t-stat=-6.87, CI = [-0.37, -0.19], df =10, P=4.32E-5; sLAN, Suppl. Figure 1L & Figure 4G, Avg. Correlation Index: -0.32 (±0.045 SE), t-test, t-stat=-7.28, CI = [-0.43, -0.23], df =10, P= 2.65E-5).

**Figure 4.**
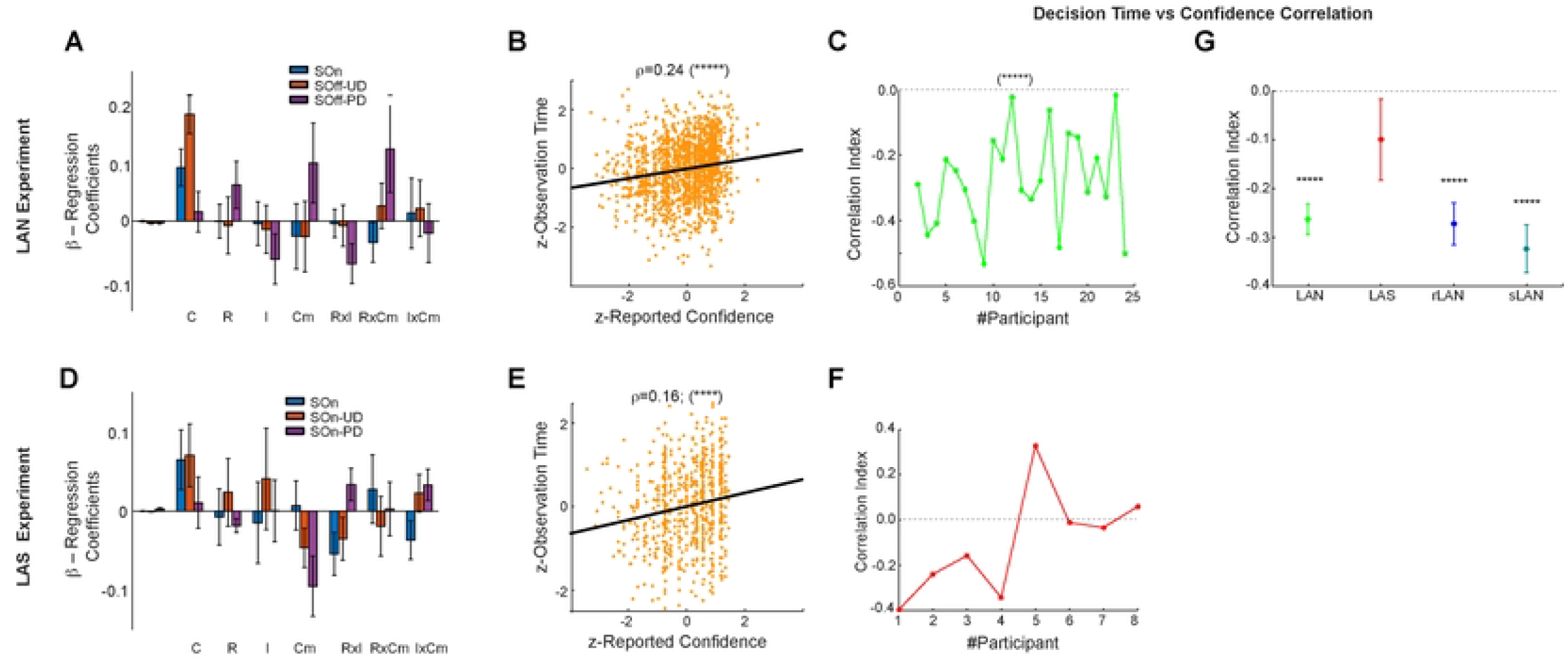
**A**. Group average and standard error β−Regression Coefficients of a GLM of the observation time of the Selected Option during the SOn, SOff-UD and SOff-PD intervals vs. the indices of Confidence (C), Reward (R), Interest (I), Complexity (Cm), plus interactions: RxI, RxCm, IxCm for the LAN experiment. Metrics are z-scored. Confidence is the only predictive regressor of the time devoted at looking at either image. **B**. Scatter plot of the observation time of the selected image until decision time vs. the reported confidence (Correlation Index r=0.24, P<0.0001). **C**. Pearson correlation index per participant of the observation time until decision time with the reported confidence index (avg. 0.16; P<0.0001). **D-F**. Same as A-C for the LAS experiment. **G**. Average and std. error correlation index of the observation time until decision time vs. the reported confidence for the four tasks.

Also, to assess the growing influence of confidence during the decision, we used a GLM regressing the proportion of observation time devoted to the image ultimately selected as a function of the reported cognitive indices (confidence, reward, complexity, interest; See Methods; Equation 2) during the Stimulus-On (SOn), Stimulus-Off Until Decision (SOff-UD) and Stimulus-Off Post-Decision (SOff-PD) intervals. Group significance tests across participants showed that confidence was the only cognitive factor marginally predicting the choice during the early SOn interval of the LAN (t-test; t-stat=1.98, CI = [-0.0065, 0.29], df=21, P= 0.06), of the LAS experiment (t-test; t-stat=2.39, CI=[0.0033, 0.19], df=8, P=0.044), and of the rLAN experiment (t-test; t-stat=-1.98, CI = [-0.16, 0.01], df=10, P=0.076). However, this influence became strongly significant during the following SOff-UD interval of the LAN (t-test; t-stat=3.024, CI = [0.040, 0.21], df=21, P=0.0062; Figure 4A), LAS experiment (t-test; t-stat=3.02, CI = [0.11, 0.30], df=8, P=0.0010; Figure 4D), and rLAN experiment (t-test; t-stat=-2.27, CI = [-0.41, -0.0041], df=10, P=0.043; Suppl. Figure 2A). Also, the relationship between observation time of the selected image and confidence during this last interval is reinforced by a complementary (Pearson) correlation analysis, yielding a r=0.24 (P<0.0001; LAN Experiment; Figure 4B), R=0.16 (R<0.0001; LAS Experiment; Figure 4E), and R=-0.45 (rLAN Experiment; Suppl. Figure 2B). The sLAN exhibited no significant effects. The remaining cognitive factors, known to influence decision-making in other situations, did not yield any significant effect in this task.

To further characterize the coupling between gazing dynamics and confidence, we assessed the number of visual alternations between images (Number of Change-of-Target fixations; #CoT), as well as their frequency (fCoT: #CoT normalized by the duration of the interval of interest). Our prior was that these metrics should increase with the difficulty to determine the appropriate choice, implying more difficulty to gain confidence. We calculated both metrics during three specific phases of the decision-making process. First, during the initial stimulus presentation (Stimulus-On); second, during the subsequent phase until the decision is reported (SOff-UD for the LAN task and SOn-UD after the GO for the LAS task and until the decision is reported); third, during the post-decision interval (SOff-PD). A first analysis of the #CoT and fCoT for the LAN and LAS tasks shows that they are higher during Stimulus-On (**LAN**: Group Avg. #CoT∼4, fCoT∼1.3Hz, Figure 5A-B; **LAS**: #CoT∼6, fCoT∼1.4Hz, Figure 5D-E), dropping down to a much slower pace when they were removed and until a decision was reported (SOff-UD, Group Avg. #CoT∼1.5, fCoT∼0.25Hz, Figure 5A-B; SOn-UD during LAS: Group Avg. #CoT∼2, fCoT∼0.25Hz, Figure 5D-E). Finally, #CoT rises back up during the post-decision interval (SOff-PD; LAN: Group Avg. #CoT∼3, Figure A-B; fCoT∼1.25Hz, Figure 5D-E; SOn-DP during LAS: Group Avg. #CoT∼3, Figure 5D-E; fCoT∼1.25Hz, Figure 5B). We considered two compatible explanations for the processes underlying these modulations. First, the initial higher fCoT during the Stimulus-On period was consistent with the low initial evidence/confidence, while the lower fCoT until afterwards during the Stimulus-Off period could suggest a slower waging process caused by the deliberative process operating in working memory alone; as if retrieving details from absent images required a stronger introspection, which in the absence of stimuli, is facilitated by the dynamics of visual saccades. Second, during the reporting period, the participant has already gathered some evidence and is more confident about which option to select, hence the steeper sigmoid slopes during Stimulus-Off (Figure 3B) and the lesser need for alternatively visual fixations between image stimuli (Figure 5). To further assess the influence of the presence/absence of the stimuli on the dynamics of gazing during decision-making, we explicitly assessed the difference between LAN and LAS group distributions of #CoT and fCoT metrics. The absence/presence of the stimuli yielded no significant difference during all three intervals of interest: for #CoT (SOn-UD interval, t-test: t-stat=-0.37, CI=[-2.15, 1.49], df=28, p=0.72; SOff-UD vs SOn-UD intervals, t-stat=0.08, CI=[-2.41, 2.63], df=29, p=0.93; SOff-PD vs SOn-PD: t-stat=0.70, CI=[-2.84, 5.82], df=29, p=0.49) and for fCoT (SOn-UD interval, t-test: t-stat=0.21, CI=1E-3*[-0.61, 0.76], df=28, p=0.83; SOff-UD vs SOn-UD intervals, t-stat=0.09, CI=1E-3*[-0.60, 0.66], df=29, p=0.93; SOff-PD vs SOn-PD: t-stat=0.70, CI=1E-3*[-0.47, 0.97], df=29, p=0.49). Finally, we regressed the confidence index as a function of the #CoT and fCoT metrics with a GLM (Eqs. 3 & 4), calculated independently for each participant, metric and interval of interest, and assessed grouped significance with a t-test across the resulting regression coefficients, for each metric and interval. This yielded a strong statistical dependence for both oculomotor dynamical metrics on confidence, on the LAN experiment during the SOff intervals, but not during the SOn (LAN: #CoT; SOn t-test, t-stat =-1.41, CI=[-0.087, 0.016], df=21, p=0.17, SOff-UD t-test, t-stat =-9.07, CI=[-0.24, -0.15], df=21, p=1E-8, SOff-PD t-test, t-stat=3.91, CI=[0.04, 0.014], df=21, P=8.13E-4; Fig 5C. Note that the statistics are very similar for #CoT and fCoT). Importantly, this analysis yielded that the relationship between the reported confidence and the #CoT and fCoT metrics was negative during SOff-UD prior to the decision, and positive after that (SOff-PD interval --- Fig 5C, F). In other words, the #CoT and fCoT prior to the decision reflect the participant’s confidence, increasing/decreasing the number of saccades between stimuli when the confidence is low/high. To further test whether the presence of the stimuli exerted an influence on the relationship between gazing dynamics and confidence, we also performed the same analyses on the data from the LAS experiment, yielding a trend consistent with the LAN effects for each interval (Figure 5F), although marginally non-significant (LAS: SOn t-test, t-stat=-1.20, CI=[-0.14, 0.044], df=7, p=0.26, SOn-UD, t-stat=-2.10, CI=[-0.22, 0.017], df=7, p=0.073, SOn-PD, t-test, t-stat=2.01, CI=[-0.01, 0.13], df=7, PD=0.083; Fig 5F). Along with the observation that the influence of confidence on gazing dynamics remained during the rLAN and sLAN controls (Suppl. Figure 4), our results are overall consistent with a cognitive process driving oculomotor movements to facilitate evidence gathering from working memory (LAN), and from the stimuli in the environment --- whenever available (LAS). In brief, these results support that oculomotor dynamics are strongly modulated by the degree of confidence until the commitment for a specific option is reported. Likewise, oculomotor dynamics for the case of this particular task are independent of the presence or absence of the stimuli.

**Figure 5.**
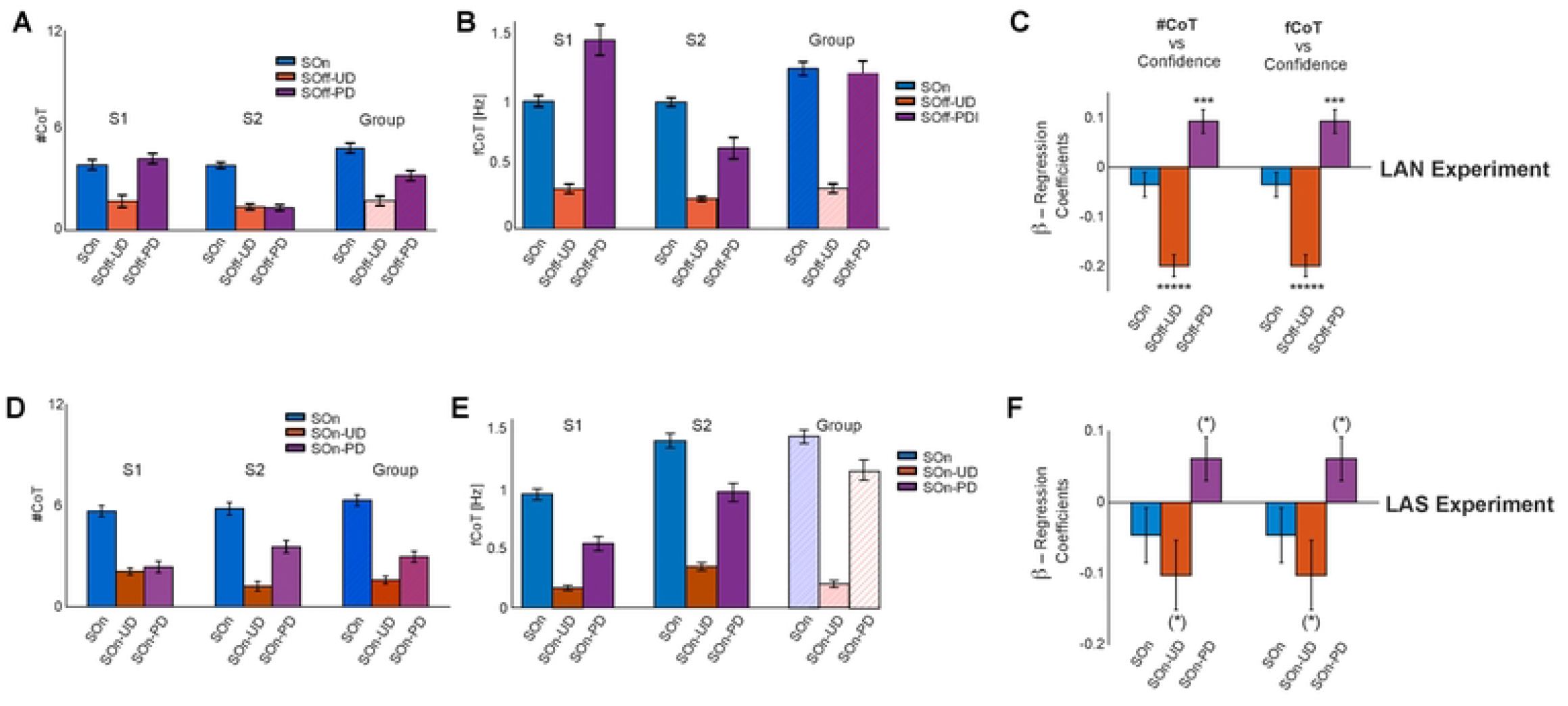
Oculomotor dynamics analyses. **A**. Average and standard error of the number of alternative fixations between the visual options (#CoT) for two typical participants (S1 and S2) and group average during the LAN Experiment, calculated during three intervals: Stimulus-On (SOn), Stimulus-Off Until Decision (SOff-UD), and Stimulus-Off Past Decision (SOff-PD). **B**. Same as A, but for the frequency of alternative fixations (fCoT) metric. **C**. Group average regression coefficients of two independent GLMs, between confidence and #CoT, and between confidence and fCoT, left and right, for the same three intervals SOn, SOff-UD and SOff-PD, during the LAN experiment. **D-F**. Same as A-C, but for the LAS Experiment during its equivalent time intervals. *Note that #CoT and fCoT, covary negatively with confidence, both when the stimuli are visible (LAS) and invisible (LAN) prior to the decision, and positively after the decision*.

Altogether, these results strongly suggest a cognitive option-assessment process that depends on oculomotor dynamics, and must therefore operate sequentially. This is supported by the increased alternation of saccades between stimuli when the options to decide upon require significant assessment. However, how sequential visual alternations are parsimonious with the parallel option representation for decisions between simpler motor actions remains to be clarified^27^. Although a complete answer to this question escapes the reach of this study, we can nonetheless qualitatively probe this by formulating and testing distinct predictions from three normative models of choice relating ocular dynamics and confidence within the context of a drift diffusion formulation of decision-making^3^, each grounded in one of three hypotheses: parallel, sequential or a hybrid consideration of options (see Suppl. Material for the full account of this formulation). We tested their predictions using the data from the four experiments described. In brief, we developed three theoretical derivations encompassing deliberation, evidence gathering and confidence, consistently with the three hypotheses as mentioned previously: in the case of parallel option consideration, the level of confidence depends on the sum of times devoted to observing each stimulus; this dependence was the difference of their inverses for the case of sequential consideration (FIG 6E). We also included a hybrid model that combined the two predictions to entertain the possibility of decision-making as a combined sequential and parallel operation. We used the confidence index reported after each decision by the participants and the observation times of each trial to test the predictions of the three models, within each of the four experiments separately. Please note that the intended purpose of using this framework is not to establish any new theory, but rather to allow a comparison between hypotheses. Ultimately, we performed a Bayesian model comparison to determine a winner model within each experiment, by inverting each of the three models via variational Bayes^28^ and obtaining a goodness of fit per participant and experiment^29^. Then, we applied Bayesian model selection to assess the model frequency and exceedance probabilities obtained from each model inversion^30^ and experiment to ascertain the hypothesis receiving the strongest support in each experiment. Figure 6A-D shows that the hybrid model is the most frequent winner in all four experiments (Model Frequency=1.0; Exceedance Probability=1.0, in all four experiments). As further validation, FIG 6F shows the average confidence index provided the participants (in quartiles) matched with the corresponding group average decision times on those trials, for each experiment (FIG 6F), along with the grand average model predictions for each experiment. Consistent with the data gathered by the previous experimental analyses, these results strongly favor the hypothesis that the dynamics of evidence accumulation in this visual task extend beyond the typical parallel dynamics observed during action selection, and incorporate elements of sequential dynamics, biased with the dynamics of visual fixations, even when there are no visible stimuli.

**Figure 6.**
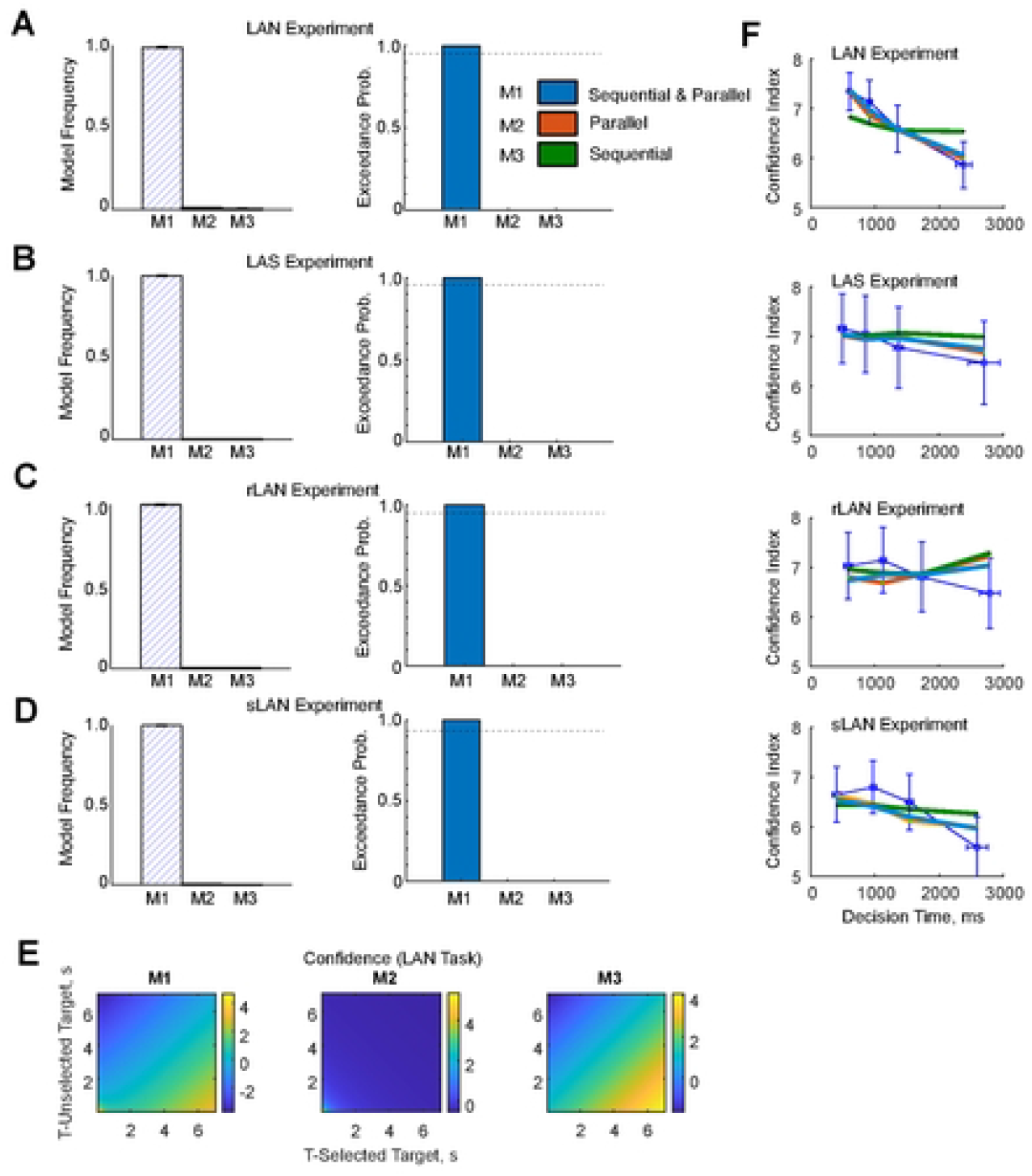
A model of mixed parallel and sequential processing of option-stimuli explains the relationship between observation time and confidence the best. **A**. Bayesian Model Comparison metrics (Model Frequency & Exceedance Probability) obtained from a Random-Effect Analysis on the three hypotheses relating confidence and observation time (sequential-parallel, sequential, parallel) for the participants of the LAN Experiment 1. **B**. Same as A, but for the LAS Experiment. **C**. Same as A, but for the rLAN Experiment. **D**. Same as A, but for the sLAN Experiment. **E**. Prediction of Confidence for all three Models on the LAN Experiment. **E**. Average Prediction of Confidence as a function of the time devoted to look at the selected vs non-selected option, based on the posteriors obtained from inverting the model for all participants of LAN experiment. **F**. Confidence across participants (quartiles) as a function of decision-time (mean and std. error; thin blue line) for all four experiment, along with the average prediction for all three models (same color coding of A-D).

## DISCUSSION

Here we studied how oculomotor dynamics relates to reported confidence, and influences decision-making in a series of tasks in which the participants had to choose the preferred of two options, presented by images on a screen. Importantly, the choices were to be made in a specific and unusual context, introduced by a phrase at the beginning of each trial. In three of the four tasks, participants had to report their choices when the images were non-visible. To characterize the influence of the presence of the options on the decision itself, we first assessed differences of decision times between LAN and LAS tasks, finding no statistically significant differences. Next, we assessed the differences in terms of frequency and duration of alternating fixations between images during a first interval in which the option images were shown (Stimulus-On) and the second reporting interval during which the images were absent (Stimulus-Off), finding no statistical differences in either of both intervals. Finally, we assessed the relationship between their reported confidence and oculomotor metrics; the fraction of time devoted to looking at either image or its related empty frame, and the number of visuo-motor alternating fixations at either image option. This yielded a significant positive correlation between observation time of the image selected and the reported confidence, and a negative one with the number of alternative fixations between option images. Finally, our computational analyses of the relationship between image observation times and confidence yield the prediction that the dynamics of decision-making combine elements of sequential sampling and parallel option deliberation, providing a path to reconcile the simultaneous consideration of motor options^31^ and the sequential sampling typical of complex, value-based decision-making^3^. Furthermore, our results also suggest an internal deliberative process during option assessment, which operates in working memory and is partially independent of the availability of sensory evidence. In this context, oculomotor dynamics play a role similar to a memory index that sequentially facilitates the processing options in a sequential fashion, as a function of the image being currently scrutinized. Moreover, the control of oculomotor dynamics is cognitively modulated as a function of the level of confidence during deliberation.

Although motor decision-making is typically described by models that assume that options are encoded and evaluated in parallel^1,32^, and a selection mechanism for a choice can be implemented by mutual inhibition between neurons that encode different options^33,34^, other models have proposed that some aspects of decision-making occur sequentially^3,10,25,35–37^. A parallel account predicts that confidence depends exclusively on the total time devoted to looking at either option, whereas a sequential model predicts that confidence depends on the difference of the times devoted to looking at each option. Our analyses show that our data matches neither of these qualitative predictions, thus supporting the idea of a mixed parallel and sequential option consideration and computation sub-processes integrating the decision-making process. More importantly, we have introduced intervals of time where the options to decide upon were invisible, creating an effective decoupling between sensory processing and internal deliberation; the fact that during LAN intervals, oculomotor patterns closely resemble those during the actual physical presentation of the option images strongly suggests that the evaluation of the options is being performed in a quasi-sequential fashion. While only neural data can resolve the question of what processes happen and when they happen during deliberation, our analysis predicts that the encoding ensues in parallel, with evaluation is most likely to be guided by an internal attentional mechanism, consistently with the mind’s eye hypothesis^38^.

Previous work has eloquently shown that gazing patterns correlate with choices when stimuli are visible^3,25^. These observations have been recently extended to memory tasks where stimuli become invisible after a presentation period. In one study where several options and their locations need to be memorized before the reporting choice stage^39^, it has been shown that oculomotor patterns are different between the visible and non-visible option conditions. It is possible that in that study, the large number of options to be remembered (six in total) overloads working memory, thus slowing down decision-making during the LAN interval due to the impeded memory retrieval. This result contrasts with those of our binary choice task: even though contexts are abstract and uncommon and options complex, participants behave no differently in the LAN than in the LAS tasks, possibly because holding two options in working memory is within reach. Our study also included confidence reports and how they relate with the oculomotor patterns, and although some work has shown that looking time and number of changes of gazed target correlates with confidence during the presentation of visual options^10,40,41^. To our knowledge, ours is the first showing that a similar pattern holds when stimuli are invisible. Again, this critical observation suggests that deliberation is in part a sequential process facilitated by gaze patterns in a situation where options are associated to locations in the visual field –the look-at-nothing effect^24^. In conclusion, our results speak in favor of a series of sequential operations to describe decision-making in the context of this task.

## Materials and Methods

### Participants

A total of forty-two participants participated in four versions of a decision-making task: Look at nothing (LAN) experiment; 24, 12F+12M, 18-35yold; Look at Something (LAS); 12, 6M+6F; 18-35yold; Motor reversal (rLAN); 12 6M+6F; 18-35yold; Short decision-time (sLAN); 10 5M+5F;18-35 yold). Their contact information was obtained through the CBC database. Participants had no known neurological damage and normal vision. Due to limitations of the eye tracker device (Eyetribe, Inc, DK), which could not maintain stable tracking of light coloured eyes or of participants wearing glasses, these had to be excluded from our study. Furthermore, since the main task was instructed in English, we required participants to hold at least a B2 level of English to participate in our study. Participants were also instructed not to wear make-up during the sessions to avoid light reflexes. Maximum session duration extended for 1.5h, receiving 5€ every 30min as monetary compensation. All participants signed an informed consent form prior to initiating the experimental session and all methods were carried out in accordance with relevant guidelines and regulations.

### Ethics Statement

Ethics regulations for this project were approved by the local ethics committee with Ref #0100. All participants were provided with an informed consent form explaining the experimental procedures, rights of the participants and data protection. This information was also verbally explained to each participant prior to the experiment. Their written consent was requested before the beginning of the experiment. All the participants were debriefed after the completion of the experiment. The information of the experimenter was provided for further questions.

### Apparatus and Stimuli

Experiments were performed in a soundproof room at our laboratory. The task apparatus consisted of a comfortable chair, placed in front of a table with a Samsung 20-inch LED monitor with 1600×900 resolution (first 12 participants of the 1^st^ experiment only) or --- thanks to lab renovation, an HP Omen 25-inch LED monitor with 1920×1080 resolution (last 12 participants of the LAN experiment and all participants of experiments LAS, rLAN, sLAN experiments). The screen was placed across the table 45cm from the participant, and was used to show the visual stimuli to the participant. An EyeTribe (EyeTribe, Inc, DK) tracking device was used to track and record eye movements and pupil size data from participants. The oculometer was placed on the table right under the screen, aligned with it and perpendicular to the participant’s line of sight. The tracker’s server version was 0.9.56, and we used a 12-screen data point routine to calibrate the system prior to starting the task. The participant was instructed to lean his/her head on a chinrest to maintain a fixed distance from the table/oculometer, to minimize their head movements during the experiment and to rest their eyes and hands during the breaks, and to hold her right hand index and middle fingers on two consecutive keys of the computer to report their choices. During the calibration process, participants were instructed not to move their heads after the calibration process, including the break times between the blocks, as to preserve the oculometer calibration profile throughout the experiment. Task flow control was performed with a Python custom-built script, implementing the Tkinter Graphical User Interface (GUI). The data from each session was transferred to a MySQL database (Oracle, CA), and analyzed with custom-built scripts in MATLAB (The Mathworks, Natick, MA), licensed to the Pompeu Fabra University.

### Procedure and Tasks

This study included four similar experimental tasks. All four were decision-making tasks in which participants were instructed to make decisions between two images shown on the screen. Each task addressed a specific aspect of the decision-making process, or control for a specific confound. The four tasks labelled as follows: 1) look-at-nothing (LAN): to specifically address the dynamics of decision-making when visual stimuli were absence, 2) Look-at-something, LAS: same as LAN but retaining the stimuli on the screen during the decision interval; 3) motor reversal (rLAN): same as LAN but inverting choice and motor response side; 4) short observation (sLAN): same as LAN but with a shorter (2s) image observation interval. Preparation for the four tasks was identical and is described next. After the task was explained to the participant and she/he was ready to start, we proceeded to calibrate the oculometer. To this end, the participant was instructed to approach the table and place the chin onto the chinrest to constrain head movements. We used the 12-point calibration profile provided by the EyeTribe software. The calibration was repeated until it reached a minimum score of 3 (out of 5). Participants not reaching a score of 3 were excluded from the experiment. Each experimental session consisted of eighty trials, equally distributed into four blocks. Each trial introduced a different context or situation in which two options were presented by two images, right and left in the screen, to decide upon. To introduce the participants to the task and familiarise themselves with the task dynamics, they performed two trials guided by the experimenter, and trained during ten additional trials.

The timeline of a typical trial of the LAN task is described next (see Figure 1). Each trial began by showing a 2×2cm black cross in the centre of the screen. After 1s, the crosswire disappeared, and a sentence describing the context of the decision was shown in the centre of the screen. After 1s, the text disappeared and a second crosswire was presented in the centre of the screen. After 1s, the Stimulus-On (SOn) interval started, showing two images related to the context were shown on the screen, left and right, equidistant from the centre of the screen, and from the top and bottom of the screen. After 4s, the Stimulus-Off (SOff) interval started with a GO, signalled by removing the images and leaving the frames behind. The SOff interval extended for 4s and was divided into two sub-intervals: from the GO Until the Decision is reported (SOff-UD), and until the end of the 4s (Post-Decision; SOff-PD). At the GO signal, the images were replaced by two empty green frames of the same size. Participants reported their choice by clicking the left/right mouse button if they select the left/right image, and with a maximum duration of 4s. After the decision was reported or after 4s, the participants were asked to quantify four cognitive aspects of their decision: complexity, interest and reward associated to the selected image, and the confidence in their decision, by rating them with a number [0-9]. Each question-answer lasted 6s. At each question, participants were given the option of jumping to the next by pressing the keyboard enter key, after reporting each rating (Figure 1). If the trial ended before a fixed trial duration of 28s, participants were required to wait until that trial duration had elapsed. We used this method to encourage participants to think carefully about the questions asked and to simultaneously provide them with a sense of control of the trial flow. The time slot for each block was 12m 42s. When participants completed all 80 trials, they were shown a message on the screen to inform them that the experiment had ended. At that point, they were free to disengage from the chinrest and the experimenter entered the room for debriefing and payment. The differences in the timeline of the four Tasks are described next. The difference between LAS and LAN tasks, is that the images were shown during the initial image presentation interval, and after the GO signal during the decision reporting as well. The rLAN task was different from the LAN in so far; the image screen side and the motor response were reversed. In other words, if the participant wished to choose the right image, he/she had to press the left mouse button and vice-versa. Finally, the difference between sLAN and LAN tasks is the shorter SOn interval (2s).

### Context and Images

We designed a total of 90 contexts, based on situations in which we would not respond automatically in our daily lives to construct imaginary scenarios where to make decisions. At the beginning of each session, the participants trained to make decisions for 10 contexts, the responses to which were excluded from further analysis. The contexts were introduced by a single sentence printed on the screen, and binary options were presented by pictures showing objects or elements fitting each context. The image resolution was 265×256 pixels. Most of the pictures were obtained from two free datasets: BossStimuli, BradyLab. A few were collected via Google image search from the websites which did not require attribution and were available for non-commercial use. The list of contexts and images are available at this respository: https://www.kaggle.com/datasets/novecentous/lanimages/data. The order of presentation of the contexts in each session was one of three different randomized orders.

### Behavioural Analyses

Analysis of the processes underlying participants’ decisions was carried out through careful examination of gaze trajectories over the screen stimuli, and keypress responses with custom written MATLAB (The Mathworks, Inc; Natick, MA) code. We used mouse button-press to detect and quantify reactions times, and carefully analysed gaze trajectories to characterize the decision-making processes. Trials were discarded if the response time was negative or longer than 4s. On a single trial basis, we calculated the decision time (DT) and, from the gaze trajectory, we extracted the amount of time spent gazing over each image presented and over their locations once these were removed from the scene. From gazing trajectories, we also detected the amount of times participants change their focus from one image to the other. We performed pair-wise comparisons between the DT distributions obtained by means of t-tests between experiments. To assess effect size we also calculated the Bayes factor associated to these tests.

### Observation Time vs Choice Likelihood

To analyse the cognitive processes of option analysis and deliberation during decision-making, we resolved to analysing eye movements and quantifying the time durations devoted to look at either stimulus, or at the frames marking their former locations. It was expected that participants will choose more often the image (and the location where the image had been formerly presented) they watch the longest. To test this for each participant, we quantified the logarithm of the ratio of the time duration devoted to look at the right over the left image for each trial, distributing them into quartiles, and calculated the number of times the participant chose the right over the left image for each quartile. We performed this calculation separately for the observation and for the reporting intervals. Second, we fitted a psychometric logistic curve to the data^42^,

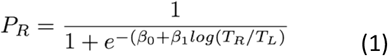

where β_0,_ β_1_ are the regression coefficients and log(T_R_/T_L_) the ratio between right/left observation time. This curve was fitted on a single participant basis, obtaining a coefficient of determination, and a p-value quantifying the goodness-of-fit and confidence intervals. To assess significance per participant, we used a permutation test. In brief, we recorded the β_1_ slope parameter obtained from the original fit. We then compared this value to the distribution of β values obtained from 10,000 shuffle data sets, in which the right choice preference values obtained for the four quartiles were randomly shuffled. If the original β_1_ was larger than 95% percentile of the distribution of β_1_ obtained from the fits of the shuffled data, the result was considered significant at P<0.05. To determine group significance, we performed a t-test on the set of β_1_ logistic regression coefficients (the sigmoid slope parameter) obtained from the original data, as a test that the distribution of slopes is significantly different from zero, establishing significance at P<0.05.

### Cognitive Factors and Observation Time

The fixation patterns indicate which stimuli are informative or interesting about each option^43^. Furthermore, eye movements can also be informative about the location of the previously encoded stimuli in the scene and the decision strategies used during memory retrieval^44–46^. Here we recorded scales of the four cognitive factors listed next, which each participant was asked to rank with a number [0-10] after reporting each decision: 1) how Confident (C) are you of your choice; 2) how Rewarding (R) was the choice; 3) how Interesting (I) was the choice; and 4) how Complex (CM) was the choice, as potential factors participating of the decision-making process. As part of our gazing strategy analysis, we fitted a Generalized Linear Model (GLM) to the time devoted to observing the selected option and the values reported for the different cognitive factors: Confidence (C), Reward (R), Interest (I), Complexity (CM), and the interactions RxI, RxCM, IxCM. The fitting was performed separately for three intervals: Stimulus-On (SOn), Stimulus-Off (SOff), divided into two intervals: from the GO signal Until the Decision is reported (SOff-UD), and Past the Decision (SOff-PD) and until the 4s SOff interval is over. In some cases, we also used aggregates of these intervals for specific analyses.

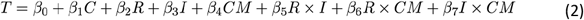

T stands for the proportion of time devoted to look at the selected option over the total interval duration (SOn, SOff-UD, SOff-PD). Group significance was assessed with a t-test over the *β*_*i*_ group coefficients obtained for each factor across participants, for each individual experiment, and established at a threshold of P<0.05.

### Deliberative Saccadic Movements and Cognitive Factors

Decisions may often favour the preferred option. However, when the difference between options is minimal and/or hard to assess, visual decisions based on a single saccade per image are bound to be uninformed. Furthermore, in our case, decisions should be based on the appropriateness of each option concerning the context, which demands some extra effort of future projection. Under these circumstances, we hypothesized that participants would use a strategy of alternately gazing to each image to gradually gather the evidence associated to each option, and to ultimately choose the most befitting option to each scenario ^25^. To assess this, we first defined two metrics: the number of times participants fixated from target to target (#Changes of Target - #CoT, which is an integer number), and the Frequency of the Changes of Target (fCoT), defined as the #CoT divided by the duration of the interval of study. We calculated these two metrics within three time intervals: Stimulus-On (SOn), Stimulus-Off Until Decision (SOff-UD) and Stimulus-Off Past-Decision (SOff-PD). Second, to quantify the relationship between these metrics and the cognitive factors presented previously, we calculated two GLMs between the z-scored metrics and the z cognitive factors reported (see EQ3-4)

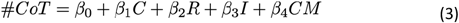

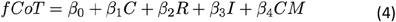

These two GLMs were fitted on the data of each participant. Group significance was established by running a t-test on the *β*_*i*_ group coefficients obtained for each factor across participants, for each individual experiment.

### Confidence and Deliberation with Parallel and Sequential Diffusion-Models

In addition to the data-based analyses described until now, we also inquired about two hypotheses by which evidence could be accumulated by a drift-diffusion model in a binary decision-making task: by maintaining simultaneous consideration of both options (parallel hypothesis), or by alternating focus between both options (sequential hypothesis). To assess which hypothesis got the most of support, we first formalized both hypotheses as two diffusion models, in which choices occur as the accumulation of evidence reaches one of two bounds^1,32^, and then performed a Bayesian model selection between them (see section next). Importantly, these two hypotheses have significantly different implications as to how we relate evidence accumulation to the confidence with which we commit to a decision, which we formalize for the case of here studied. The goal of this exercise was to obtain defining features for each model that relate observation times and confidence, and then perform a model comparison of the extracted features, as described next.

We based our analyses on the participants reported confidence after each decision they made. The common dynamics of both models may be described as follows. We assume that each option has an associated value μ_1_ and μ_2_, and that these are unknown to the participant. The preferred option is determined by the estimated difference between those values, which is determined by gathering noisy information associated to each option for some time, to ultimately choose the option with the highest value. Furthermore, although the participant may get estimates for either option (1 or 2), their associated noise renders the choices necessarily uncertain, and the probability of choosing the most valuable choice smaller than one, which has been formally assimilated to a quantification of the notion of confidence^37,47^.

In the ***parallel diffusion model***, the dynamics of accumulated evidence x(t)at time t obey to

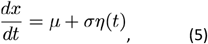

where μ = μ_1_ ― μ_2_ is the drift, σ^2^ is the variance, and η(t)is a zero-mean, unit-variance white noise process. The drift represents the average instantaneous value difference between the two options, which is corrupted by noise, turning the drift estimation into an inference problem. The noisy evidence in the right-hand side of Eq. (5) is accumulated over time as x(t), starting from zero. In this model, the accumulation of evidence occurs in parallel for both options --- in the form of the relative difference between them. The accumulation of evidence stops when x(t)hits for the first time one of the decision boundaries θ or –θ, which determines both the choice and the decision time. The option 1 is chosen if the upper bound is hit first; x(t)= θ, as then the probability that μ = μ_1_ ― μ_2_ > 0 is larger than one half. The option 2 is chosen if the lower bound is hit first; x(t)= ―θ.

By assuming a flat normal prior over value, the (Bayesian) decision confidence g(t)at decision time t in such a model may be characterized as

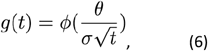

where *ϕ* is the cumulative density function of the standard normal^47^. In this context, the decision confidence equals the probability of the participant being correct in the decision given that the accumulated evidence is x(t)= θ at decision time t^4,47^. Remarkably, the previous expression is exact and applies even if the bounds depend on time θ(t), in any arbitrary fashion. In other words, decision confidence depends exclusively on decision time, and not on the unknown drifts, assuming all other parameters of the model being fixed. Therefore, **longer decision times will involve a lower decision confidence** regardless of the individual fractions of time looking at each option, confirming the intuition that if long times are required to make a decision, it must have been because the choice was difficult and the drift was close to zero.

By contrast to the parallel hypothesis and model just described, in *the* ***sequential diffusion model*** a single option is attended at any given time t^3^. This we can model by means of a diffusion model, where the accumulated evidence x (t)obeys to equation 5, with the difference that the drift μ(t)is now time-dependent. This drift reflects the average instantaneous evidence for the option that is currently attended, and equals μ(t)= μ_1_ when that is option 1, and μ (t)= ―μ_2_ when that is option 2. Thus, in this model, the accumulation of evidence occurs sequentially, alternating between options 1 and 2, according to the time devoted to observing each of them. As in the parallel model, we assume that there are two time-independent decision boundaries, and that the choice and the decision time is determined by a time at which one of the boundaries is hit as a result of the accumulated evidence.

Under specific assumptions, we may also derive an expression for decision confidence for the sequential model as well, which corresponds to the probability of choosing correctly. To that end, we assume that the options are of the same absolute value but of opposite sign (an offset can be added without modification of the equations), but the participant does not know whether μ_1_ = ― μ_2_ = μ_0_/2 or μ_2_ = ― μ_1_ = μ_0_/2, that is, it is not known which of the two options is better. Therefore, the goal of the participant is to determine whether μ_1_ = μ_0_ or μ_1_ = ―μ_0_. It can do so by alternatively attending to each of the options during specific time durations t_1,1_,t_2,1_,t_1,2_… where t_i,j_ is the time devoted to observing option i = {1,2} during an epoch j. The sufficient statistics for this problem are simply the accumulated evidence Δx_1_ and Δx_2_ during the total times Δt_1_ = ∑_j_ t_1,j_ and Δt_2_ = ∑_j_ t_2,j_ that options 1 and 2 were paid attention to, respectively^47^, and then the probability that μ_1_ = μ_0_ is

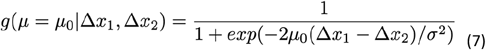

This probability is larger than one half only if Δx_1_ > Δx_2_, that is, when the evidence while observing option 1 rises above the evidence while observing option 2. This case supports the hypothesis that the first option is better than the second one. Finally, assuming the values are small, we have that Δx_1_ ≈ μ_0_t_1_ and Δx_2_ ≈ μ_0_t_2_. After inserting them into Eq. (7) we obtain

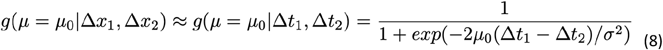

From equation 8 follows that decision confidence in the **sequential diffusion model** does not depend on the total decision time t = Δt_1_ + Δt_2_, but on their difference ΔT = Δt_1_ ― Δt_2_. Specifically, a longer observation time of option 1 implies a higher confidence that this option will exhibit the highest value.

These derivations are based on a specific mathematical formulation of confidence in the context of the drift diffusion model (Please, see ref. 44 for the full analysis). The reasoning hereby introduced is merely intended to allow a qualitative comparison of the parallel vs sequential operation of oculomotor dynamics and confidence, in the terms described in the experiment. In this light, we formulated two strictly different hypotheses: parallel (Eq. 9) and sequential (Eq. 10). However, it is also possible that evidence accumulation abides to a compromise between both models. Because of this, we also propose a third mixed sequential and parallel model (Eq. 11), where the accumulation of evidence is a combination of both. In this case, the decision confidence (*C in formulae 9-11*) will correlate with both total decision time and the difference of observation times between them. These hypotheses introduced lead to the following models:

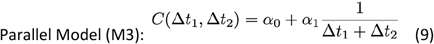

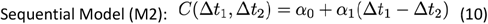

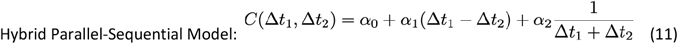

The comparison procedure is described next.

### Bayesian Model Selection

We used Bayesian Model Selection (BMS) to assess which formulation of evidence accumulation best explained the decision times obtained experimentally and the confidence reported by each participant after each trial, according to the hypotheses/models described in the previous section. Comparison between hypotheses was performed as a function of the individual free energy, by means of a Random Effects Analysis (RFX) fitting the [0-10] confidence index reported by the participants, normalized between 0 and 1. We performed Bayesian Model Selection (BMS) across the RFXs at the group level^30^. Briefly, models are treated as random effects, which could differ per participant, though assuming a fixed distribution across the population. To make the test possible, we first needed to use Variational Bayes (VB)^48^ to obtain the posteriors and a frequency metric for each model (M1-M3, cf. eqs. 9-11) across the population^28,28,49^. The advantage of the VB algorithm is that it does not only invert models with a robust parameter estimation, but also estimates the model evidence, representing a trade-off between accuracy (goodness-of-fit) and complexity (degrees of freedom)^50^. The free energies estimated for each participant and model were submitted to a random effect analysis (RFX), which assumes that models could differ between participants and that they have a fixed (unknown) distribution across the population.

The BMS procedure yielded the exceedance probability^30^, which measures how likely it is that any given model is more frequent than all other models in the comparison set. The BMS yields one additional metric, the model frequencies (MF) across that population, which explains the larger amount of variance for each participant’s data. The specific model comparison was based on the distribution of PFs, assumed to be Gaussian, characterized by two parameters: a mean and a standard deviation. We performed the BMS procedure on the three aforementioned models.

## List of Acronyms

Abbreviation: Definition
LAN: Look-At-Nothing Experiment
LAS: Look-At-Something Experiment
rLAN: Motor Reverse Look-At-Nothing Experiment
sLAN: Short Observation Look-At-Nothing Experiment
#CoT: Number of Changes of Target
fCoT: Frequency of Changes of Target
UD: Until Decision
PD: Post Decision
TR: Target Right
TL: Target Left
C: Confidence
R: Reward
I: Interest
CM: Complexity

## Acknowledgments

IC was supported by the European Union’s Horizon 2020 research and innovation programme/Human Brain Project (Marie Sklodowska-Curie Research Grant Scheme IF-656262), by the Spanish Project PID2019-105093GB-I00 (MINECO/FEDER, UE) and CERCA Programme of the Catalan Government, and by the European Union’s Horizon 2020 Framework Programme for Research and Innovation under the Specific Grant Agreements No. 945539 (Human Brain Project SGA3). RM was supported by the Howard Hughes Medical Institute (HHMI; ref 55008742), The Bial Foundation (106/2022), Ministerio de Ciencia e Innovación (Ref: PID2020-114196GB-I00/AEI) and ICREA Academia.

## Author contributions

Conceptualization: IC, GS, PM, RM

Methodology: IC, GS, PM, RM

Investigation: IC, GS, PM, RM Visualization: IC, GS

Funding acquisition: RM, IC

Project administration: RM,

IC Supervision: RM, PM

Writing – original draft: IC

Writing – review & editing: IC, RM, PM

## Competing interests

Authors declare that they have no competing interests.

## Data and materials availability

All data, code, and materials used in the analysis are available at this repository: https://www.kaggle.com/datasets/novecentous/lanimages/data.

## Notes

### Competing Interest Statement

The authors have declared no competing interest.

